# A drastic shift in the energetic landscape of toothed whale sperm cells

**DOI:** 10.1101/2021.03.01.432277

**Authors:** L. Q. Alves, R. Ruivo, R. Valente, M. M. Fonseca, A. M. Machado, S. Plön, N. Monteiro, D. García-Parraga, S. Ruiz-Díaz, M.L. Sánchez-Calabuig, A. Gutiérrez-Adán, L. Filipe C. Castro

## Abstract

Mammalia spermatozoa are a notable example of energetic compartmentalization. While mitochondrial oxidative phosphorylation is restricted to the midpiece, sperm-specific glycolysis operates in the flagellum. Consequently, these highly specialized cells exhibit a clear adaptability to fuel substrates. This plasticity is essential to ensure sperm motility, and is known to vary among species. Here we describe an extreme example of spermatozoa-energetics adaptation. We show that toothed whales exhibit impaired sperm glycolysis, due to gene and exon erosion, and demonstrate that dolphin spermatozoa motility depends uniquely on endogenous fatty acid *β*-oxidation, but not carbohydrates. Our findings substantiate the observation of large mitochondria in spermatozoa, possibly boosting ATP production from endogenous fatty acids. This unique energetic rewiring emphasizes the physiological body reorganisation imposed by the carbohydrate-depleted marine environment.

## Main Text

The rise of modern human societies from ancient civilizations implied a division of labor and specialization: critical leverages towards the evolution of complex interactions. Such trend is also noticeable in the evolution of biological complexity, associated with the emergence of structured compartments and resource distribution towards specific functions (e.g. multicellularity). At the cellular level, we can also recognize structural partitions. Yet, the compartmentalization of metabolic pathways, in time and space, within such structures is less intuitive (*1*). Mammalian spermatozoa provide an illustrative example. The energy in the form of ATP production, vital for motility, capacitation, and fertilization, is subcellularly separated in sperm cells. While glycolysis provides a local, rapid, and low-yielding input of ATP along the flagellum fibrous sheath, oxidative phosphorylation (OXPHOS), far more efficient over a longer time frame, is concentrated in the midpiece mitochondria (*2*). The relative weight of glycolysis and OXPHOS pathways in sperm function is species-dependent, and sensitive to oxygen and substrate availability (i.e., glucose, fructose, fatty acids, lactate, glycerol, ketone bodies, or pyruvate) (*3*). For example, while OXPHOS was suggested to be predominant in bull spermatozoa, in humans and mice, glycolysis is essential, notably in the distal tail (*4, 5*). Besides partitioning energy production, sperm cell energetics display an additional singularity: the occurrence of sperm-specific gene duplicates and alternative spliced variants, although with conserved function (Fig. S1) (*6*).

The wider selective forces driving the compartmentalization and adaptability of this energy system in mammalian species remain largely unknown; much like the impact of ecosystem resource availability (e.g. carbohydrates, fatty acids, proteins) and dietary adaptations in reproductive physiology traits (*7*). Here, we investigated the Cetacea, an iconic group of fully aquatic and carnivorous marine mammals, evolutionarily related to extant terrestrial herbivores. In this lineage, episodes of profound trait adaptation have been accompanied by clear genomic signatures (*8-10*). To investigate the evolution of sperm energetics, we started by analyzing the glycolysis rate-limiting enzyme, glyceraldehyde-3-phosphate dehydrogenase, spermatogenic (*Gapdhs*) (*11*), in mammalian genomes. Using a pseudogene inference pipeline, *Pseudochecker* (*12*), we scrutinized the coding condition of *Gapdhs* in 163 genomes and found that this gene displays a *Pseudoindex*, a probability measure of pseudogenization (functional inactivation) (*12*), compatible with gene lesion events in an extremely restricted number of mammalian lineages (Table S1; table S2) (*13*).

In effect, this pattern is almost exclusively limited to toothed whales (Odontoceti, Cetacea). Next, we employed manual validation (Fig. S2) to inspect the coding sequence of *Gapdhs* in species with a *Pseudoindex* higher than 2, including 18 species of toothed whales (*13*), unveiling numerous gene-inactivating mutations in Odontoceti species (Fig. 1A; fig. S3-S21). Further examination of a bottlenose dolphin (*Tursiops truncatus*) sperm transcriptome showed expression of *Gapdhs* with an aberrant splicing pattern, with sequence reads encompassing the identified genome mutations (*13*) (fig. S22). A single mutation is shared across all the examined Odontoceti species, with exception of the early diverging beaked (Kogiidae) and sperm (Physeteridae) whales, which display independent inactivating mutations (Fig. 1A). *Gapdhs* ORF-disrupting mutations were also identified in other mammals, but were only found to be valid disrupting mutations in the naked mole-rat, known to have simplified and degenerate sperm (*14*), and Pholidota (pangolins) (fig. S4, S23-27). In addition, we identified relaxed selection in the whole Cetartiodactyla clade, including baleen whales, Pholidota, Carnivora, and Cingulata clades (Fig. 1B; table S3). Since we could not exclude the possibility of a functional substitution by the somatic paralogue of *Gapdhs* (*Gapdh*), we next investigated *Pgk2*, a glycolytic isozyme expressed uniquely during spermatogenesis and responsible for the first ATP-generating step in this pathway (Fig. S1) (*15*). Despite overall synteny conservation of the genomic *locus* in cow, hippopotamus, and baleen whales, together with no significant selection relaxation or intensification (Table S3), we found that *Pgk2* is sequence-deleted in all examined toothed whales (Fig. 1C; fig. S28-29). Again, *Pgk2* inactivating mutations were also deduced in pangolins and the naked mole rat (Fig. S29-32). Out of the ten enzymes responsible for the conversion of glucose to pyruvate in sperm cells (Fig. S1), we established that the sperm-cell restricted genes *Gapdhs* and *Pgk2* display severe sequence lesions in Odontoceti species. Importantly, besides sperm-specific genes, other glycolytic isoenzymes arise from alternative splicing, securing an independent regulation from somatic splice variants (*16*). That is the case for Aldolase A (*AldoA*), responsible for the synthesis of glyceraldehyde-3-phosphate, and presenting a testis-specific splice variant, *Aldoa_v2*, with a distinct N-terminus exon (*6*). Thus, we next examined the sequence integrity of this sperm-exonic variant (Fig. 1D). We first established that *Aldoa_v2* emerged in the ancestor of placental mammals and is expressed uniquely in testis (Fig. 1D). Moreover, we deduced the presence of ORF-disrupting mutations within the sperm-specific exon in all of the examined Cetacea species (Fig. 1D, fig. S33-60).

**Fig 1.**
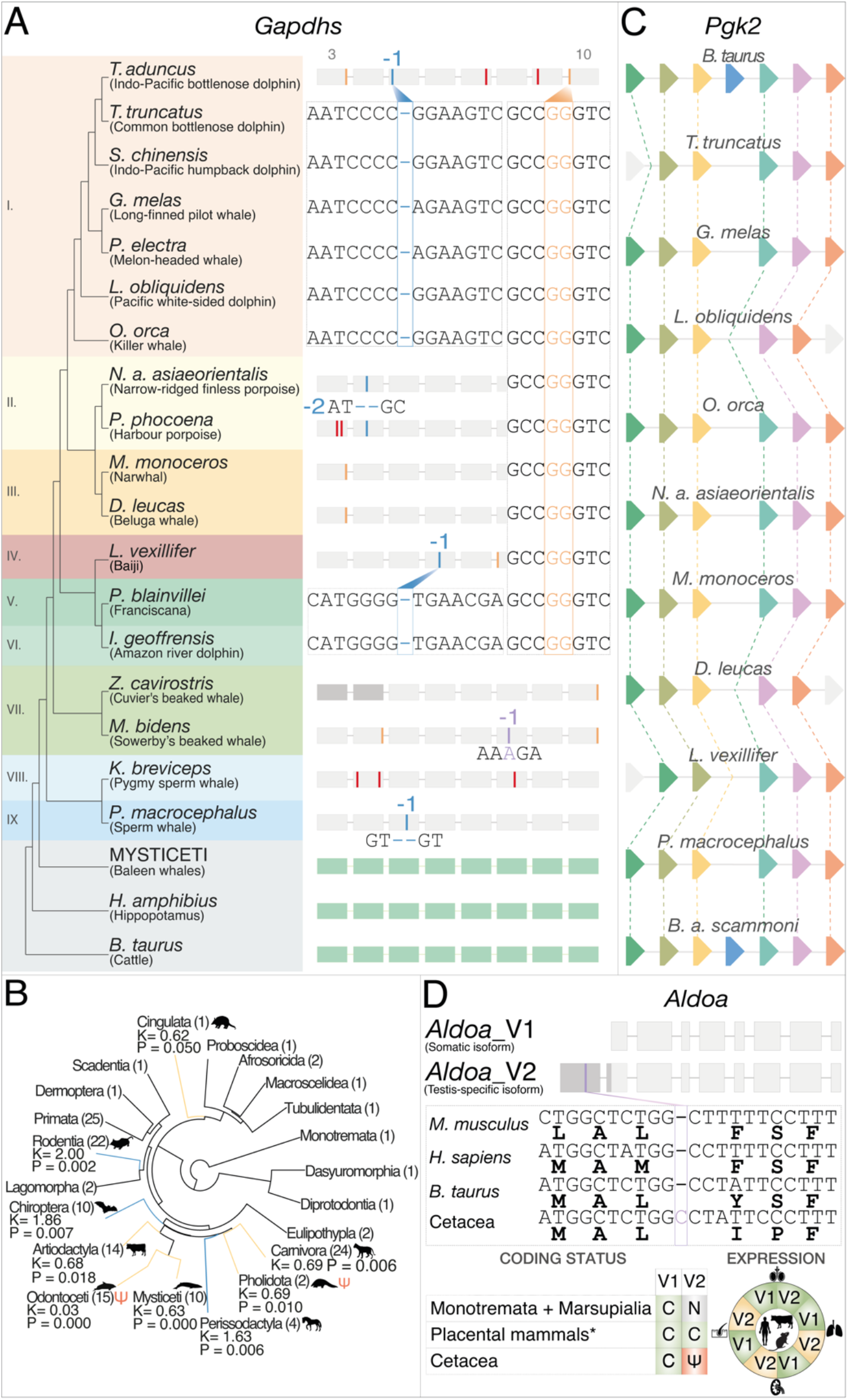
Erosion of sperm-specific glycolysis genes. (**A**) The mutational landscape of *Gapdhs* in Cetacea and hippopotamus (*H. amphibius*). Examples of the detected gene disruptive mutations (insertions in purple, deletions in blue, in-frame premature stop codons in red, and splice site mutations in yellow) are represented according to their position within the affected exons (numerated on top, from the 3rd to the 10th exon – full representation available in fig. S4). Odontoceti families are represented as follows: I. Delphinidae, II. Phocoenidae, III. Monodontidae, IV. Lipotidae, V. Pontoporiidae, VI. Iniidae, VII. Ziphiidae, VIII. Kogiidae, IX. Physeteridae. (**B**) Branch-specific relaxation parameters inferred for *Gapdhs* in mammals. (**C**) Comparative synteny maps of the *Pgk2* genomic locus in Cetacea and *Bos taurus* (cow). Orthologous genes are joined by lines (fig. S28). (**D**) *Aldoa_v2* exon erosion in Cetacea. C-Coding, N-Not found, Ψ-Pseudogenized, Green-Expressed, Yellow-Not expressed. Silhouette images were retrieved from http://phylopic.org/.

A fully operational glycolysis pathway converts glycolysable sugars to pyruvate, with a net gain of 2 ATPs. Sugar alcohols, such as sorbitol and glycerol can also be metabolized entering the glycolytic pathway (*5*). The resulting pyruvate, at the intersection of glycolysis and mitochondrial ATP production, can be (*a*) reversibly metabolized to lactate, generating NAD^+^ which can reintegrate the glycolytic pathway, sustaining additional ATP production (Fig. 2); or (*b*) transferred to the mitochondria, metabolized to acetyl-CoA and incorporated in the Krebs cycle and OXPHOS, generating up to 36 ATP molecules (Fig. 2). Additionally, acetyl-CoA may originate from glycolysis-independent sources such as fatty acid oxidation or ketone body catabolism. Together, most of these metabolic steps are controlled by spermatozoa-restricted gene orthologues, including lactate dehydrogenase, *Ldhc*, which oxidases NADH (*17*); a pyruvate specific transporter, *Mpc1l* (*18*); a converter of pyruvate to acetyl-CoA performed by the pyruvate dehydrogenase complex composed of several genes, including the sperm restricted *Pdha2* (*19*) (Fig. 2); glycerol kinase 2 (*Gk2*), which catalyzes the production of glycerol-3-phosphate from glycerol, which can be enzymatically converted into glyceraldehyde-3-phosphate (*20*) (Fig. 2); and, the 3-Oxoacid CoA-transferase 2 (*Oxct2*) which converts ketone body-derived acetoacetate into acetoacetyl-CoA in sperm cells (*21*) (Fig. 2). We examined each gene orthologue sequence in various mammal species and found numerous ORF-disrupting mutations or exon-loss events in most Cetacea species (Fig. 2; fig. S61-S158). Overall, our results indicate the full dismantling of key energy-producing pathways in Odontoceti species: including glycolysis, pyruvate and lactate production, mitochondrial pyruvate uptake and conversion to acetyl-CoA and mitochondrial ketone body catabolism in sperm cell. The latter agrees with the previously reported inactivation of ketone body synthesis in Cetacea (*22*). In contrast, Mysticeti exhibit sparse gene loss events, with the inactivation of ketone body usage and pyruvate to acetyl-CoA conversion, suggesting a metabolic dissociation between glycolysis and mitochondrial pathways. Yet, in Mysticeti, glycolysis is most likely not fully abolished. The erosion of *Aldoa_v2* and retention of a functional *Gapdhs* suggest that partial glycolysis could occur from non-sugar substrates with the production of the intermediary glyceraldehyde-3-phosphate from glycerol; also, *Aldoa_v2* could be functionally substituted by the somatic paralogue. This landscape of gene erosion suggests a profound restructuring of energy production in Odontoceti sperm cells, and to a lesser degree in Mysticeti. Together, these findings raise the hypothesis of OXPHOS as the sole energy provider in sperm cells, a physiological arrangement previously unknown in mammalian spermatozoa. Yet, a critical question emerges: How do Odontoceti sperm cells obtain the required acetyl-CoA to feed mitochondrial OXPHOS? A natural candidate is mitochondrial fatty acid *β*-oxidation.

**Fig 2.**
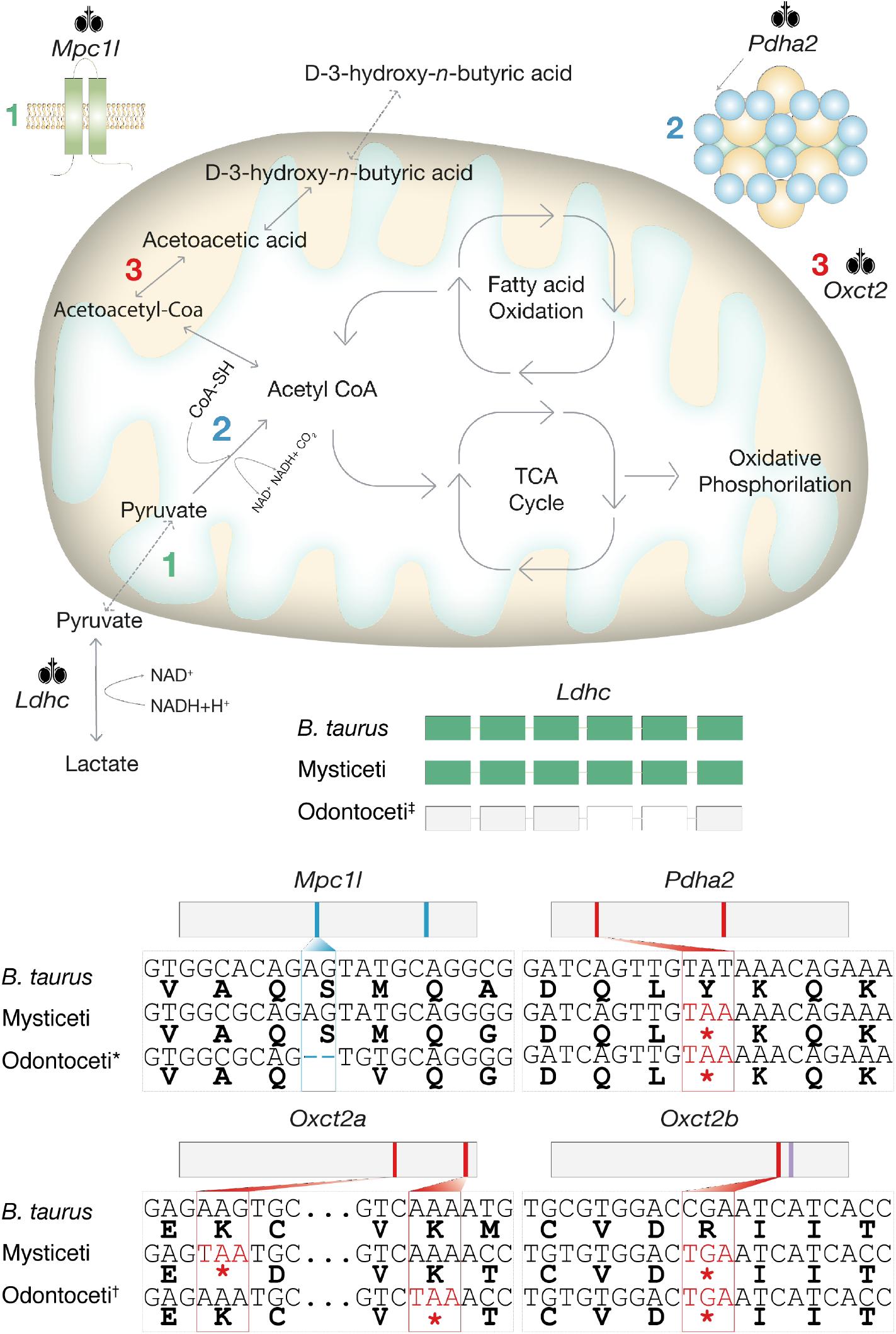
Metabolic reactions and their sperm-specific genes participate in the mitochondrial cellular respiration: oxidative decarboxylation of pyruvate, ketone body catabolism, fatty acid β-oxidation, citric acid cycle, and oxidative phosphorylation. ORF-disrupted genes in Cetacea are highlighted together with their location in the pathway. Multiple sequence alignment of the coding sequences for *Bos taurus* (cow), Mysticeti (baleen whales), and Odontoceti (toothed whales) emphasize the presence of disruptive mutations or exon loss (*Ldhc*, white rectangles). Nucleotide deletions are represented in blue, insertions in purple, and in-frame premature stop codons in red. *except *I. geoffrensis* (Amazon river dolphin), *P. blainvillei* (La Plata dolphin), Ziphiidae (beaked whales), *P. macrocephalus* (sperm whale), and *K. breviceps* (pigmy sperm whale). †except *P. macrocephalus* (sperm whale) for *Oxct2a* and *O. orca* (killer whale) for *Oxct2b*.

To experimentally validate this hypothesis, we examined dolphin sperm motility and movement longevity, an indicator of exhaustion of reserve energy sources, in the presence/absence of the glycolytic substrate glucose, the glycolysis end-product pyruvate, as well as etomoxir, a specific inhibitor of carnitine palmitoyl-transferase 1a (*Cpt1a*) and blocker of mitochondrial fatty acid *β*-oxidation (Fig. 3A-D) (*23*). Dolphin sperm cells maintained normal motility for more than 3 days, even without glucose and pyruvate supplementation (Fig. 3A and 3B), in clear contrast with mice and bull sperm cells, which typically stop moving or display reduced mobility shortly after 1 and 3 hours, respectively (*23*). Either with or without supplementation, the percentage of cells exhibiting progressive or irregular motility was similar, with only a minute increase in hyperactivated motility when glucose and pyruvate were added to the incubation media (Fig. 3A and 3B). In contrast, when exposed to etomoxir, cell motility dropped after only 1 hour of incubation, significantly differing from the control [Control 1h: 65.3% ± 4.4% (mean ± S.E.); etomoxir 1h: 24.8% ± 2.8%, (p < 0.01)], with progressive motility practically disappearing after 2h of incubation (Fig 3C and D). An increased number of spermatozoa with hyperactive-like motility was also observed, when compared to the control group, at 1 and 2 hours of incubation with etomoxir (Fig 3D). This hyperactive-like state is different from normal hyperactive motility (displaying an increased amplitude of lateral head displacement and decreased straight-line and curvilinear velocities), but similar to that observed in subpopulations of bull sperm that die very quickly after freezing/thawing (*24*). At 24 hours of incubation, only a small percentage of sperm cells (9.3% ± 1.4%) remained motile, displaying irregular motility patterns (Fig 3D). Etomoxir-dependent *β*-oxidation inhibition was previously shown to decrease sperm motility in various species including humans (*25*) or boar (*26*). Yet, in dolphin sperm, *such* inhibition yielded faster and more pronounced effects on sperm motility, with progressive motility diverted to hyperactive-like states, ultimately leading to irregular sperm movement patterns. Overall, these findings showcase the complete dismantling of the glycolytic pathway in sperm cells while support the central role of *β*-oxidation in ATP generation to fuel normal sperm motility.

**Fig 3.**
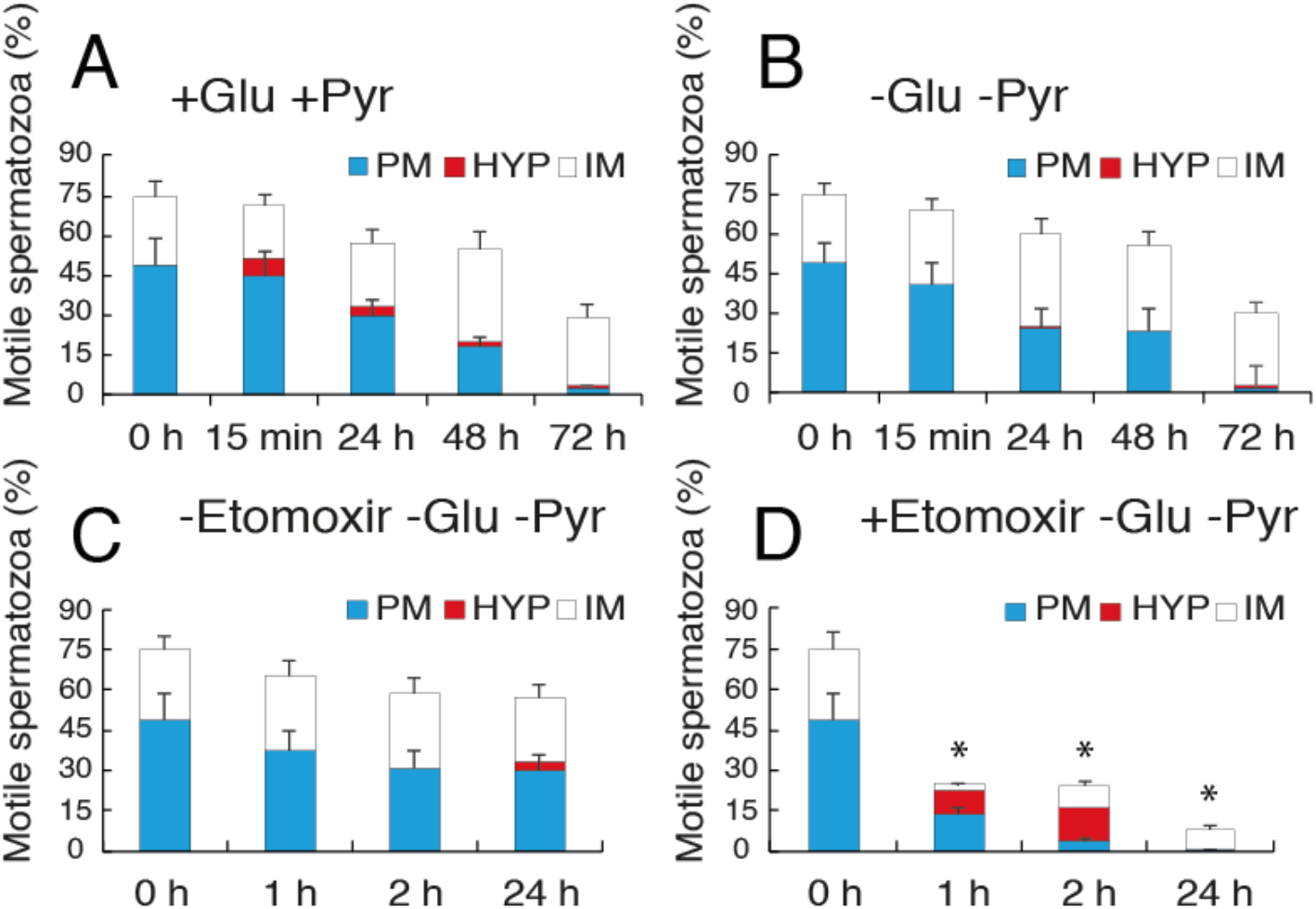
Energetic substrate assays of dolphin sperm motility. Dolphin sperm samples were incubated in the presence (**A**) or absence (**B**) of glucose and pyruvate. Total, progressive (PM; Movie S1), hyperactive (HYP; Movie S2), and irregular (IM; Movie S3) motility measured using the Integrated Semen Analysis System (ISAS) at 15 min, 24 h, 48 h, and 72 h. No differences in total or progressive sperm motility were observed between (A) and (C). Sperm samples were incubated in the absence (**C**) or presence (**D**) of etomoxir; motility was measured using ISAS at 1 h, 2 h, and 24 h. Total and progressive motility dropped significantly in the presence of etomoxir. Data are expressed as mean ± S.E. \**P* < 0.01.

To further identify the endogenous metabolites able to sustain the described cellular behavior, we compared sperm from bottlenose dolphins and bull using proton nuclear magnetic resonance (^1^H NMR) spectroscopy. A total of 12 metabolites were identified in the lipophilic phase and 16 metabolites in the hydrophilic phase (Fig. 4A and 4B). As expected, fatty acid content, in the form of free fatty acids, triglycerides, or cholesterol esters, was significantly higher in dolphin sperm, supporting the use of these lipophilic compounds as substrates for mitochondrial energy production (Fig. 4A). Phospholipids, notably phosphatidylethanolamine, were also more abundant in dolphin sperm, as well as free cholesterol (Fig. 4A). In the hydrophilic phase, L-acetylcarnitine, citrate, creatine, and glycerol content were higher in dolphin sperm than in bull (Fig. 4B). L-acetylcarnitine is the acetylated and most common derivative of L-carnitine, required for *Cpt1a*-dependent fatty acid shuttling into mitochondria for *β*-oxidation; citrate is an intermediary of the Krebs cycle, resulting from the condensation of oxaloacetate and acetyl-CoA, and a feedback inhibitor of glycolysis; creatine participates in the creatine-phosphocreatine shuttle, an enzymatic energy transport system relying on phosphocreatine diffusion from the mitochondria and subsequent regeneration of ATP and creatine in the cytoplasm (*27*); and glycerol a major component of triglycerides and phospholipids. Together these findings support a drastic modification of sperm energetics in Odontoceti species towards fatty acid *β*-oxidation-fueled OXPHOS.

**Fig 4.**
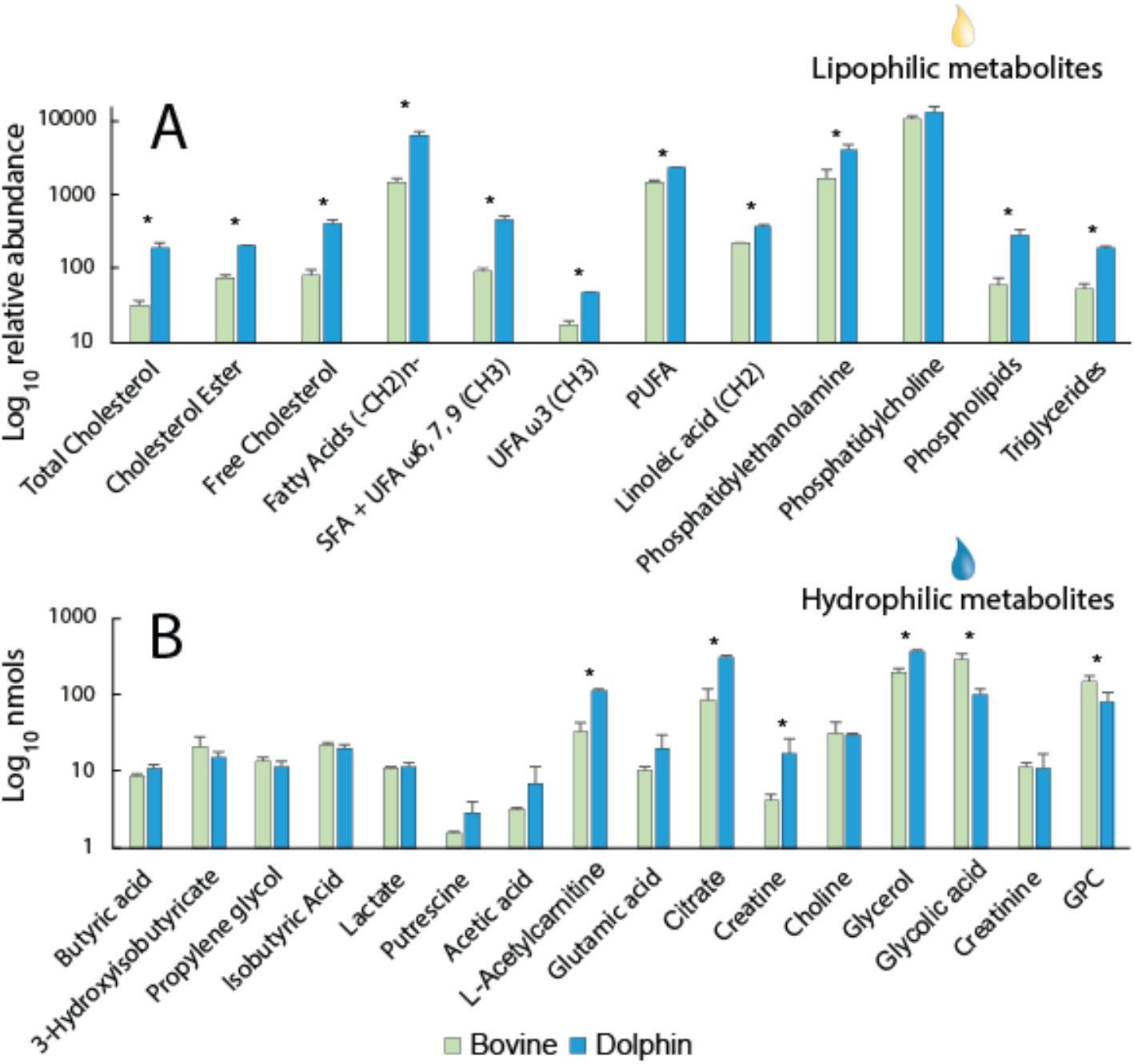
Metabolites identified in bull and dolphin sperm extracts by proton nuclear magnetic resonance (^1^H NMR) spectroscopy using lipophilic (A) and hydrophilic (B) phases. \**P* < 0.01. FA: fatty acid; SFA: saturated fatty acid; UFA: unsaturated fatty acid; PUFA: polyunsaturated fatty acid; GPC: glycerophosphocholine.

Unlike other mammals, the complete demolishing of glycolysis and shift towards mitochondrial-enclosed metabolic pathways allowed the maintenance of normal sperm functions in Odontoceti. For instance, the inactivation of glycolytic genes, in humans and mice, induces defects in sperm structure and motility leading to infertility in humans and mice (*28-31*) (Table S4); while in the eusocial naked mole-rat, with a dominant male mating strategy, sperm is abnormal (*14*). Yet, in dolphins, sperm quality is generally very high, few abnormal sperm, in agreement with their promiscuous mating system based on sperm competition (*32*). This unique metabolic rearrangement was possibly accompanied by an increase in mitochondrial activity and/or ATP diffusion along the flagellum (i.e. creatine-phosphocreatine shuttle). Curiously, such metabolic burden on mitochondria is paralleled by an also peculiar phenotype in Odontoceti when compared to other Artiodactyla, including Mysticeti: bigger mitochondria with augmented cristae, resulting in larger midpiece volumes (*32*) (Fig S159 and S160).

The evolution of such a unique energetic system in Odontoceti sperm cells most probably emerged as a consequence of dietary adaptations and resource availability (*33*). Odontoceti dietary intake, such as that of the common bottlenose dolphin, is characterized by high-protein and low-carbohydrate content, leading to an overall low glycolytic activity (*34*). Glucose to fatty acid shifts in energy metabolism have also been described in Cetacea skeletal muscle as a result of gene inactivation (*9*). Surprisingly, circulating glucose levels in healthy dolphins are high (*35*). The fate of such high glucose stores is most likely the brain. Brain metabolic energy is exclusively provided by glucose or fatty acid-derived ketone bodies; yet, the synthesis of the latter is impaired in Cetacea (*22*). Odontoceti, but not Mysticeti, have large brain-to-mass ratios and encephalization quotients similar to non-human primates (*36*). Such critical concatenation of events likely diverted the usage of glucose, a scarce resource in aquatic environments, from peripheral functions in order to support the elevated energy demands of the brain (*37*). Additionally, this energetic system, yielding good quality and enduring spermatozoa, is probably favored in a polyandry system with sperm competition, such as that exhibited by many Odontoceti species, and facilitates cell motility in complex vaginal folds (or pseudocervices) hypothesized to restrict sperm access to the upper reproductive tract (*38*). Our findings also have vital implications for captive reproduction programs across the world. Considering that dolphin sperm utilizes fatty acids as an energy source supporting motility, the formulation of sperm storage media may need to be revisited.

## Acknowledgments

This research was funded by COMPETE 2020, Portugal 2020, by the European Union through the ERDF, grant number 031342, by FCT through national funds (PTDC/CTA-AMB/31342/2017), and by grant RTI2018-093548-BI00 from the Spanish Ministry of Science and Innovation. R.V. is funded by the FCT under the grant SFRH/BD/144786/2019. The study was also supported by Trainers and veterinarians of Oceanogràfic Valencia, especially Carlos Barros-García, for the obtaining and transfer of dolphin ejaculates. S.R.D. is supported by a “Doctorados Industriales 2018” fellowship of Comunidad de Madrid (IND2018/BIO-9610).

## Author Contributions

Conceptualization, L.F.C.C.; methodology, L.Q.A, R.R., M.M.F A.M.M., and A.G.-A., L.F.C.C; validation, L.Q.A, R.R., R.V., and A.G.-A., L.F.C.C.; formal analysis, L.Q.A, R.R., R.V., M.M.F., A.M.M., N.M., S.P., D.G.-P., S.R.-D., NM.J.S-C., A.G.-A. and L.F.C.C.; investigation, L.Q.A, R.R., R.V., M.M.F., A.M.M., N.M., S.P., D.G.-P., S.R.-D., M.J.S-C., A.G.-A., and L.F.C.C.; resources, J.G.-A. and L.F.C.C.; data curation, L.Q.A, R.R., R.V., M.M.F., A.M.M., N.M., S.P., D.G.-P., S.R.-D., M.J.S-C., A.G.-A., and L.F.C.C. writing-original draft preparation, L.Q.A., R.R., A.G.-A. and L.F.C.C. writing-review, and editing and supplementary Materials writing, L.Q.A, R.R., R.V., M.M.F., A.M.M., N.M., S.P., D.G.-P., S.R.-D., M.J.S-C., A.G.-A., and L.F.C.C; supervision, L.F.C.C. project administration, L.F.C.C. funding acquisition, A.G-A., and L.F.C.C. All authors approved the submitted manuscript. Competing interests: The authors have no competing interests to declare. Data availability: The clean RNA-seq read datasets were submitted to the Sequence Read Archive (SRA) database of NCBI, and can be consulted under the Bio project number (PRJNA703781).

## Supplementary Materials

### Materials and Methods

#### Assessing the coding status of the in-study genes in Mammalia – general approach

To investigate the coding status of each target gene, the following method was employed: (*i*) Orthology of Cetacea genes was assessed combining phylogenetics using the Ensembl.org Gene tree pipeline and synteny analyses to authenticate one-to-one orthologues, and to define genomic regions to be collected for gene annotation; (*ii*) Pseudo*Checker* (pseudochecker.ciimar.up.pt) *(12, 39)* analysis and/or manual annotation of the open reading frame (ORF) of each gene was performed over previously retrieved genomic regions, with these further being inspected for gene inactivating mutations (frameshift mutations, premature stop codons and splice site mutations); (*iii*) finally, for each gene and each species, at least one identified ORF-abolishing mutation was further validated using independent genomic sequencing read sets obtained from the National Center of Biotechnology Information (NCBI) sequence read archive (SRA) database, when available. The sequence analysis pipeline is summarized in Fig. S2. Overall, a total of 169 mammalian genomes, including 27 Cetacea, were analyzed in the study (Table S1).

#### Synteny analyzes

Synteny maps were built for each of the analyzed genes in Cetacea (Figs. S3, S28, S33, S61, S68, S87, S115, S133, and S146) using NCBI-available annotated genome assemblies: *Balaenoptera acutorostrata scammoni* (minke whale: accession ID -GCF_000493695.1), *Delphinapterus leucas* (beluga whale: GCF_002288925.2), *Globicephala mela*s (long-finned pilot whale: GCF_006547405.1), *Lagenorhynchus obliquidens* (Pacific white-sided dolphin: GCF_003676395.1), *Lipotes vexillifer* (baiji: GCF_000442215.1), *Monodon monoceros* (narwhal: GCF_005190385.1), *Neophocaena asiaeorientalis asiaeorientalis* (narrow-ridged finless porpoise: GCF_003031525.1), *Orcinus orca* (killer whale: GCF_000331955.1), *Physeter macrocephalus* (sperm whale: GCF_002837175.2) and *Tursiops truncatus* (bottlenose dolphin: GCF_001922835.1).

For each target gene, the *Bos taurus* (cow) genome was screened (GCF_002263795.1), and the corresponding ortholog gene *locus* was set as a reference for Cetacea. Finally, for each studied gene: (*i*) genome assemblies with annotation of the orthologous gene, five (characterized, with available gene description or available full name) protein-coding genes, upstream and downstream of the annotated target gene were collected and placed into the syntenic map; (*ii*) concerning genome assemblies displaying the absence of annotation for the target gene, the corresponding *B. taurus* (cow) direct flanking genes were set as reference genes, further collecting the immediately four remaining upstream or downstream adjacent genes to these.

#### Gene annotation

NCBI gene annotations for the mammalian orthologues of *Gapdhs, Pgk2, Gk2*, and *Ldhc* were sequence validated *via* Pseudo*Checker* (pseudochecker.ciimar.up.pt), which estimates the coding status of a gene *(12, 39)*. To this effect, a total of 4 Pseudo*Checker* analyzes were run, using the human gene orthologue as a comparative coding sequence input (NCBI Accession ID regarding human *Gapdhs:* NM_014364.5; *Pgk2*: NM_138733.5; *Gk2*: NM_033214.3; *Ldhc*: NM_002301.5), as well as the genomic sequence encompassing the putative ORF of the orthologous gene of each target species, directly exported from the NCBI genome browser (default parameters were used, except for the minimum exon alignment identity, set to 40%). Through *PseudoIndex*, a user assistant metric built into the software, we rapidly estimated the erosion condition of the tested gene on a discrete scale ranging from 0 (coding) to 5 (pseudogenized) *(12, 39)*, we further classified the predicted sequences into functional (*PseudoIndex* between 0 and 2) or putatively pseudogenized (*PseudoIndex* higher than 2), with subsequent manual annotation and validation of eventual gene disrupting mutations (see below) (Fig. S2; Tables S2 and S5-S7).

In the case of Cetacea and *Hippopotamus amphibius* (hippopotamus), sequence annotation was performed on both annotated and unannotated genomes for all in-study genes (*Gapdhs, Pgk2, Gk2, Ldhc, Aldoa_v2, Mpc1l, Pdha2, Oxct2a*, and *Oxct2b*) (Fig. S2). In detail, (*i*) for genomes displaying annotation for the target genes, the genomic portions underlying the respective annotation were directly collected through the NCBI gene browser; (*ii*) for annotated genomes, but without the target gene annotation (e.g. *Ldhc, L. obliquidens -*Pacific white-sided dolphin), according to the previously produced synteny maps, the genomic sequences ranging from the upstream to the downstream flanking genes were collected and sequence analyzed; (*iii*) finally, for unannotated cetacean genomes (i.e., *Balaenoptera bonaerensis*, the Antarctic minke whale), the genomic sequences were collected via BLAST searches against the corresponding genome assembly using, as a query, a set of three sequences, including the *B. taurus* (cow) target gene corresponding ortholog coding sequence (CDS), as well as the CDS’s of the flanking genes in the same species. The returned BLAST hits were analyzed, further retrieving the genomic sequence corresponding to the consensus hit across those obtained per each query sequence. For the cases where no consensual blast hit was obtained, all hits corresponding to the *B. taurus* (cow) CDS query were inspected, the aligning regions submitted to a back-blast search against the nucleotide (nt) database of NCBI, with the matching genomic sequence(s) corresponding to the gene of interest being the one(s) selected for annotation (when existing) (Table S8).

The collected genomic sequences harboring the putative ORF of the in-study genes were imported into Geneious Prime 2020 software (www.geneious.com) *(40)* and the gene CDS was manually annotated using, as a reference, the *B. taurus* (cow) corresponding orthologs concerning cetaceans and *H. amphibius* (hippopotamus) and, for non-cetacean mammals, the *H. sapiens* (human) counterparts, except for *Aldoa_v2* which made use of the *Mus musculus* (mouse) reference ortholog (Table S8). Explicitly, per gene and species, by using the Geneious *(40)* built-in map to reference tool, and with the highest sensitivity parameter selected, each (3’ and 5’ untranslated region-flanked) reference species’ coding-exon (in the cases of *Gk2, Pgk2, Mpc1l, Pdha2, Oxct2a*, and *Oxct2b* genes, the corresponding reference single coding-exon) was mapped against the target genomic sequence. Due to alignment issues regarding the exon 2 of *B. taurus* (cow) *Gapdhs* against Cetacea, this was first mapped against the corresponding *H. amphibius* (hippopotamus) genomic sequence, further using the predicted exon as the reference *Gapdhs* exon 2 for annotation in cetaceans. Regarding *Oxct2a* and *Oxct2b* gene annotations, to ensure that we were not mistakenly annotating each of these for the corresponding copy, we first mapped the exons of the respective *B. taurus* (cow) flanking genes, forcing the cow ortholog of the *Oxct2* copy of interest to align within the corresponding flanked genomic sequence (*Pabpc4* -XM_010803664.2 and *Macf1* -NM_001143860.2 for *Oxct2a*; *Trit1* -NM_001192840.1 and *Ppie* -NM_001098161.1 for *Oxct2b*). The cases for which we could not map at least one of each copy’s flanking genes were rendered inconclusive. Finally, each exon alignment was further screened for gene disruptive mutations, including in-frame premature stop codons, frameshift, and splice site mutations (any deviation from the consensus donor splice site GT/GC or the consensus acceptor splice site AG) (Figs. S4, S29, S34, S62, S69, S88, S116, S134, and S147).

#### Validation of gene erosion lesions

To ensure that the identified genetic lesions were not the result of sequencing and/or genome assembly artifacts, we further validated at least one of these via the mapping of unassembled genomic sequencing reads retrieved from two independent genomic projects from the NCBI sequence read archive, when available. To accomplish this, blastn searches were conducted against the selected SRA projects, using as a query the nucleotide sequence containing the selected mutation(s). The matching sequencing reads were imported into Geneious Prime 2020 *(40)* software and mapped against the manually annotated mutation using the built-in map to reference tool (highest sensibility parameter selected), further confirming, or not, the presence of the identified mutation (Figs. S5-S21, S23-S27, S30-S32, S35-S60, S63-S67, S70-S86, S89-114, S117-132, S135-145, and S148-158).

#### Assessing the Aldoa somatic and testis-specific isoforms expression status in other mammals

To confirm the testis-restrictive expression of the *Aldoa_v2* isoform in *H. sapiens* (human), *B. taurus* (cow), and *M. musculus* (mouse), we inspected the transcriptomes of four different tissues in each of these species, including lung, kidney, skin, and testis. To accomplish this: (*i*) for each species, we have directly imported the genomic portion corresponding to the *Aldoa* gene using the NCBI genome browser into Geneious *(40)* (human corresponding genomic accession ID: NC_000016.10; cow: NC_037352.1; mouse: NC_000073.7) and, with the built-in map to reference tool (high sensitivity parameter selected), we annotated both isoforms using the mouse ortholog as a reference (*Aldoa_v1* accession ID: NM_007438.4; *Aldoa_v2*: NM_001177307.1); (*ii*) per tissue and species, we inspected two independent NCBI SRA transcriptomic projects or samples, and run a blastn search against each of them using, as reference, the respective CDS of both *Aldoa_v1* and v2 isoforms (-max_target_seqs 5000) (Table S9); (*iii*) the corresponding reads were imported into Geneious *(40)* and these were further mapped against each species respective isoform annotations (maximum read gap size adjusted to the length of each isoform with a maximum mismatch rate of 2%); (*iv*) finally, aligning reads were count per species, tissue and isoform, with isoforms displaying at least 30 aligning reads being considered as expressed (Table S9).

#### Selection analysis and RELAX

To determine the direction of natural selection (relaxed or intensified) in *Gapdhs, Pgk2, Gk2*, and *Ldhc* genes across the mammalian tree, we used the RELAX software on the Hyphy package *(41, 42)*. For this purpose, in mammal species displaying ORF-abolishing mutations in the coding region of these target genes, we manually recovered the gene sequence, removed frameshift insertions and in-frame stop codons were recorded as missing data (NNN). For species for which no evidence of disruption was found during the Pseudo*Checker* analyses, we directly collected the respective CDSs from the NCBI database. Excluding cetaceans, we considered a species as having the target gene mutated only for the cases where we could validate ORF-disruptive mutations using data from at least 1 SRA genomic project (mandatorily different from the reference genome corresponding project) or 2 distinct individuals. Mammals displaying issues related to exon alignment and fragmented genomic regions in the target gene and/or exon deletions were excluded from the analysis. In Tables S10, S11, S12, and S13, we present the list of mammals used in RELAX analysis for each target gene.

For each gene, mammalian sequences were translated-aligned in Geneious Prime 2020 *(40)* using the Blosum62 substitution matrix. RELAX estimates a relaxation/intensification parameter k that either indicates relaxed selection (k < 1) or intensified selection (k > 1). Using previous alignments as input, we ran the test by specifying (*i*) different cetacean families as foreground branches; (*ii*) each mammalian clade (or species) presenting ORF-disrupting mutations as test set; (*iii*) different mammalian orders whose number of species used in this analysis was higher than 10. For all the tests, all the other branches leading to species that did not lose the target gene were specified as background. Specifically, for *Gapdhs*, given the consensual loss pattern observed in Odontoceti, we ran RELAX considering (in contrast to other genes) (*i*) all Odontoceti species as foreground branches; (*ii*) all Mysticeti species as the test set; (*iii*) and all mammalian orders except Marsupial orders.We report all values of k and three omega classes, together with the percentage of sites under these dN/dS rates for each run in Table S3.

#### Mitochondrial size measurements across Mammalia

To further examine the mitochondrial size of mammalian sperm, we collected published photographs of longitudinal sections through the midpiece of the sperm flagellum *(43-60)*. For well-defined mitochondrion, for which we were able to see its boundaries, we took two measurements: one called “*Height*” defined as the measurement of the mitochondrion parallel to the axoneme, and another called “*Width*” defined as the measurement of the mitochondrion perpendicular to the axoneme (Table S14). Images for which only magnification was displayed, the height and width of each mitochondrion were calculated from distances measured with a ruler (± 0.05 cm). On the other hand, for images displaying scale, we used the ImageJ software *(61)* to automatically calculate both measures. For the majority of the species, data on mitochondrial size was only taken from one individual/figure (Table S14).

Finally, a scatterplot was constructed using R software *(62)*, allowing the comparison between mitochondrial sizes in different mammalian lineages (Fig. S159). As both measurements were not normally distributed, we tested for differences between mammalian orders using the nonparametric Kruskall-Wallis test. Moreover, for each measurement, each order was compared to all the others (i.e., base-mean) to determine if the height/width of their mitochondria was, significantly bigger or smaller. Boxplots were developed allowing for a visual comparison of the differences in the distribution of data across several mammalian orders (Fig. S160).

#### Ethics of experimentation

Dolphin spermatozoa were collected from two adult bottlenose dolphins (*T. truncatus*) as part of the routine husbandry training methods for the bottlenose dolphin on a voluntary basis at Oceanogràfic-Valencia (Valencia, Spain). All procedures were performed using methods consistent with the Animal Care Protocol of the Oceanogràfic-Valencia. All animals used in this project were cared for using procedures that are consistent with the regulations and policies of Oceanogràfic-Valencia and the Spanish Council on Animal Care Documents, ‘Guide to the care and Use of Experimental Animals’, ‘Categories of Invasiveness in Animal Experiments’ and ‘Ethics of Animal Experimentation’.

#### Semen collection and cryopreservation

Semen collection has been previously described *(63)*. After collection, semen samples (avoiding contact with any source of light) were diluted at a 1:1 ratio using a commercial extender [Beltsville Thawing Solution (BTS)] (GVP, Zoitech Lab, Madrid, Spain) and frozen in TRIS egg yolk-based buffer containing glycerol to obtain a 3% final glycerol concentration and a final concentration of 200 x 106 sperm/mL *(63)*.

#### Dolphin sperm RNA extraction, construction, processing, and analysis

The total RNA content was extracted using one sample of Dolphin sperm as follows. Semen samples were washed with sperm preparation medium (CooperSurgical, Denmark) and gently centrifuged at 4°C. Cell pellets were ressuspended in NZYol reagent (Nzytech, Portugal) and incubated at 65°C for 5 minutes. Following extraction procedure and done in according to the manufacturer’s instructions in combination with the Illustra RNAspin Mini RNA Isolation Kit (GE Healthcare, UK) with on-column DNase I digestion and elution in RNase-free water. The total RNA concentration was measured in a microplate spectrophotometer with Take3™ Micro-Volume Plate (BioTeK, USA) and its quality was checked through the measurement of the OD260/280 ratio values (1.8-2.0). The RNA integrity was verified by running 1μl in a 1% agarose gel. The sample was shipped to Macrogen (Seoul, Korea), the strand-specific library was built (Insert size: 250–300 bp) and the sequencing was performed with an Illumina NovaSeq 6000 platform (Paired-end 150 bp).

The RNA-Seq dataset was processed with the protocol and parameters applied in Machado et al. (2020) *(64)*. Briefly, the FastQC *(65)* (0.11.8) software was used to check the initial quality of the dataset, Trimmomatic *(66)* (0.38) to trim and quality-filter the raw reads, Rcorrector *(67)* (1.0.3) to correct the sequencing errors, and Centrifuge *(68)* (1.0.3-beta) to taxonomically classify and filter out exogenous raw reads and possible sources of contamination. All reads corresponding to match hits out of the Mammalia class (Taxon ID: 40674) were removed. To complement the dataset assessment, the clean reads were mapped onto the NCBI-available genome assembly of *T. truncatus* (bottlenose dolphin) (GCF_001922835.1) resorting to Hisat2 *(69)* (2.2.0) (default settings), calculating the rate of mapped reads with samtools *(70)* (1.9.0). The stats of raw and clean datasets, as well as the rate mapping against the reference, can be consulted in Table S15. The clean datasets were submitted to the NCBI SRA database and can be consulted under the BioProject -PRJNA703781.

Finally, resorting to the produced transcriptomic dataset, we inspected the occurrence of expression reads from *Gapdhs, Gk2*, and 5 highly testis-expressed genes according to the Human Protein Atlas (www.proteinatlas.org): *Ybx1, Phf7, Tsacc, Tnp1*, and *Eef1g*. To accomplish this: (*i*) we collected the *T. truncatus* (bottlenose dolphin) CDS of the previously annotated *Gapdhs, Gk2*, as well as the control genes respective CDS from NCBI, and ran a blast search of each of them against the clean reads dataset (-task megablast, -max_target_seqs 20000); (*ii*) the reads corresponding to each blast search were collected and mapped against the respective gene using the Geneious *(40)* built-in map to reference tool (maximum read gap size adjusted to the length of each gene with a maximum mismatch rate of 2%); (*iii*) the splicing patterns were inspected and the aligning regions covering mutated regions screened for mutated transcripts (Fig. S22). A similar approach was employed to deduce the presence of mRNA of these genes (here, also including *Pgk2)* in mouse sperm cells (NCBI SRA -BioProject -PRJNA320896).

#### Effect of glucose and pyruvate on sperm motility

Sperm starvation was done in a HEPES modified Toyoda-Yokoyama-Hosi medium (HEPES-TYH) *(71)* according to changes in the composition that have been previously indicated *(23)*, for which glucose and pyruvate were omitted. This medium can be supplemented with HCO3^-^ or BSA to support capacitation. Thawing was done in a water bath at 37 °C for 50 s. Then spermatozoa were separated on a density gradient BoviPure™ (Nidacon, Sweden) and resuspended in HEPES-TYH medium, and then washed by centrifugation for 5 min at 300g, and resuspended in 1 ml of HEPES-TYH medium. Sperm were incubated with or without glucose and pyruvate at 5% CO2 at 38°C in TYH medium supplemented with 15 mM HCO3^-^ and 5 mg/mL bovine serum albumin (BSA, fatty acid-free, A0281) for 3 days. Motility and kinetic were reported every day by the Integrated Semen Analysis System (ISAS; Projectes i ServeisR + D S.L., Valencia, Spain) and acrosome status and vitality were analyzed.

#### Fatty acid oxidation inhibition

To ascertain what energy source is used by dolphin spermatozoa, we incubated sperm samples with etomoxir, a selective inhibitor of CPT1, the rate-limiting enzyme of b-oxidation. Sperm samples from the two adult dolphins (n =6, triplicate experiment) were incubated in HEPES-TYH supplemented with 3% (w/v) fatty-acid-free bovine serum albumin and 1% (v/v) penicillin-streptomycin-neomycin at 38 °C, 5% CO2, in the presence or absence (control) of 1 mM of etomoxir. Motility and kinetic (ISAS analysis), and acrosome status were reported after 1 h, 2 h, and 18h.

#### Sperm motility and kinetics

Sperm motility was objectively determined using ISAS. Ten microliters of sperm suspension were placed in a Mackler chamber on the stage heated to 37 °C of a Nikon Eclipse E400 (Nikon, Tokyo, Japan) fitted with a digital camera, Basler acA1300-200uc (Basler AG, Ahrensburg, Germany). Three to five movies of 1.5 s were recorded at 60 frames/s using the software Pylon Viewer provided by Basler, capturing at least 100 moving spermatozoa *(72)*. The motility and sperm kinetics were analyzed using the free software ImageJ *(61)* with the plugin ISAS bmg following instructions for analyzing dolphin spermatozoa *(73)*. The parameters analyzed were as described by Mortimer et al. *(74)*: straight-line velocity (VSL; µm/s), curvilinear velocity (VCL; µm/s), average path velocity (VAP; µm/s), linearity (LIN) (defined as (VSL/VCL) × 100), straightness (STR) (defined as (VSL/ VAP) × 100), wobble (WOB) (defined as (VAP/VCL) × 100), amplitude of lateral head (ALH) displacement (µm), and beat-cross frequency (BCF; Hz). Also, we examined the percentage of spermatozoa showing more signs of hyperactivation (HYP) by determining out of all the analyzed spermatozoa (9340) the lower VCL and ALH values of the 10% of spermatozoa with the highest VCL and ALH. These values were: VCL = 150 µm/s and ALH = 5.5 µm. Thus, we defined spermatozoa showing hyperactive-like motility as those showing VCL > 150 µm/s and ALH > 5.5 µm *(75)*.

#### Sperm preparation and purification

Semen samples from 2 dolphins (*T. truncatus*) and 3 Friesian bulls (*B. taurus*) were analyzed. Frozen sperm from 2 adult bottlenose dolphins spermatozoa were separated as previously indicated on a density gradient BoviPure™ (Nidacon, Sweden) and resuspended in PBS medium, and then washed two times by centrifugation for 5 min at 300g. Frozen semen straws (0.25 mL) from 3 Friesian bulls previously tested for IVF were thawed at 37 °C in a water bath for 1 min and centrifuged for 10 min at 300 g through a gradient of 1 mL of 40% and 1 mL of 80% Bovipure (Nidacon Laboratories AB, Göthenborg, Sweden), according to the manufacturer’s instructions. The sperm pellet was isolated and washed two times in PBS by centrifugation at 300 g for 5 min. For all the samples, after the last centrifugation cells were frozen with liquid nitrogen and stored at -80 °C for further metabolite extraction.

#### Metabolite extraction for by proton nuclear magnetic resonance spectroscopy (^1^H NMR)

A significant number of cells (∼200 million) were used to extract enough metabolites for each ^1^H NMR spectroscopy experiment. Two different pools of samples with identical phenotypes were prepared: 3 bulls and 2 dolphins. Frozen seminal samples from three different Asturian Valley bulls (*B. taurus*) were used. Bulls were housed at the Cenero AI Centre [Regional Service of Agrifood Research and Development (SERIDA), Gijón, Spain], complying with all European Union regulations for animal husbandry. Additional approval from an ethical committee to conduct this study was not required. Animals were selected based on being good breeders by having AI outcomes using frozen samples. Sperm from the two dolphins used in the motility experiments were used. Thawed semen was washed by density gradient centrifugation by placing it on top of a BoviPure gradient (Nidacon Laboratories AB, Göthenborg, Sweden) of 1 mL at 80% (lower layer) and 1 mL at 40% (upper layer) and centrifuging for 5 min at 300xg. Sperm cells were then washed three times with cold PBS and cell pellets were frozen with liquid nitrogen and stored at -80 °C for further metabolite extraction. Extraction of low-molecular-weight metabolites was performed following a previously published procedure *(76)*. For the preparation of the ^1^H NMR samples, semen cell pellets were pre-treated. A methanol extraction was performed with the following protocol: samples were defrosted to room temperature for 5 minutes slowly within the ice. 1.3 mL of CHCl3:MeOH:ddH2O in a ratio 41.7:35.6:32.7 (v/v/v) was added to the sperm sample. The Eppendorf with the extraction was placed at 4°C with agitation for 4h. The mixture was centrifuged at 4°C at max-speed (∼ 25000-30000 g) for 30 mins. The upper phase of the dissolution (Hydrophilic phase) was transferred to a new 2mL Eppendorf. The down phase of the dissolution (Lipophilic phase) was transferred to a different Eppendorf too. All the samples were then dried in a Speed-Vac. For the preparation of the hydrophilic samples to be acquired in the ^1^H NMR we have resuspended the lyophilized product with 500 μL of deuterium oxide (D2O) and 0.11 μM of DSS (Sodium trimethylsilylpropanesulfonate). Samples were briefly vortexed and 500 μL of the semen extract was finally pipetted into a 5 mm ^1^H NMR tube. Lipophilic extracts were resuspended in 300 μl of DMSO-d6 with 4 mM of TPP (triphenylphosphine) and transferred to a Shigemi tube. In all cases, sample preparation was manually done at 298 K.

#### ^1^H NMR measurements

All ^1^H NMR experiments were recorded at 298 K on a Bruker 600 MHz (12 T) Avance III spectrometer equipped with a BBO (BB, 1H) probe head. For each sample, a 1D 1 H p3919gp with water signals suppression using a binomial 3-9-19 pulse with echo gradient pair (21 min). Data analysis was done using the TopSpin 3.5 software (Bruker Biospin GmbH) *(77)*. Free induction decays were multiplied by an exponential function equivalent to 0.3 Hz line-broadening before applying Fourier transform. All transformed spectra were corrected for phase and baseline distortions and referenced to the DSS singlet at 0 ppm. The hydrophilic and lipophilic samples were recorded.

The identification of the metabolites was done on the sample with the higher concentration of sperm extracts. DSS was used as an internal reference that can be used to quantify the number of different molecules. TSP shows a peak at 0 ppm, which corresponds to 9 equivalent protons. Therefore, this peak is equivalent to a 1 mM proton intensity. The peaks are then integrated and referenced to DSS. On the other hand, 4 mM of TPP was added to the lipophilic phase. Considering the contribution of 15H the peak integral corresponds to 60 mM. However, the quantification of the lipophilic phase is impossible due to the different contributions coming from the sample type of compound. For example, a signal coming phosphatidylcholine will be composed of the contribution of the -CH2-groups in their hydrophobic tail, and the length of the tail will vary randomly. Thus, the integrals can be compared as relative integrals, not absolute quantification.

#### Statistical analysis

Statistical analysis was carried out using the software package GraphPad Prism 8.0.2 for Windows (GraphPad Software, San Diego, CA, USA). Results are expressed as means ± standard error of the mean (SEM). Means were compared and analyzed using a one-tailed paired-sample Student’s t-test or repeated measures one-way analysis of variance (ANOVA), followed by Tukey’s post hoc test. Significance was set at p < 0.05.

